# A Computational Growth and Remodeling Framework for Adaptive and Maladaptive Pulmonary Arterial Hemodynamics

**DOI:** 10.1101/2023.04.20.537714

**Authors:** Jason M. Szafron, Weiguang Yang, Jeffrey A. Feinstein, Marlene Rabinovitch, Alison L. Marsden

## Abstract

Hemodynamic loading is known to contribute to the development and progression of pulmonary arterial hypertension (PAH). This loading drives changes in mechanobiological stimuli that affect cellular phenotypes and lead to pulmonary vascular remodeling. Computational models have been used to simulate mechanobiological metrics of interest, such as wall shear stress, at single time points for PAH patients. However, there is a need for new approaches that simulate disease evolution to allow for prediction of long-term outcomes. In this work, we develop a framework that models the pulmonary arterial tree through adaptive and maladaptive responses to mechanical and biological perturbations. We coupled a constrained mixture theory-based growth and remodeling framework for the vessel wall with a morphometric tree representation of the pulmonary arterial vasculature. We show that non-uniform mechanical behavior is important to establish the homeostatic state of the pulmonary arterial tree, and that hemodynamic feedback is essential for simulating disease time courses. We also employed a series of maladaptive constitutive models, such as smooth muscle hyperproliferation and stiffening, to identify critical contributors to development of PAH phenotypes. Together, these simulations demonstrate an important step towards predicting changes in metrics of clinical interest for PAH patients and simulating potential treatment approaches.

## 1 Introduction

Pulmonary arterial hypertension (PAH) is a deadly disease with numerous underlying etiologies. While many origins remain poorly understood, all forms of PAH are accompanied by characteristic hemodynamic and morphologic changes. Over the course of the disease, increases in pulmonary arterial pressure (PAP) and pulmonary vascular resistance (PVR) due to arterial remodeling subject the right ventricle to increased afterload, which can eventually cause heart failure and death (Abman et al., 2015). Understanding the time course of these hemodynamic changes and their link to specific mechanisms of arterial remodeling is key to identifying new treatment strategies, since current treatment agents, such as vasodilators, often become ineffective in late-stage disease (Ivy, 2016).

Blood vessels are highly sensitive to loads imparted by hemodynamic forces. Indeed, alterations to baseline hemodynamics of animal models in the absence of genetic abnormalities can induce PAH alone (Zhang et al., 2018; Kameny et al., 2019). Wall shear stress (WSS) caused by the frictional force of blood acting on the vessel lumen leads to changes in gene expression and phenotype of the endothelial cells lining the vessel’s inner surface. These changes can modify activation of transcription factors, such as KLF2 and KLF4, and cause endothelial-to-mesenchymal transition that drives abnormal proliferation and intimal thickening (Moonen et al., 2015; Moonen et al., 2022). Additionally, intramural stress (IMS) resulting from pressure-induced stretch in the vessel wall acts on smooth muscle cells in the media, altering their production of extracellular matrix and active tone (Humphrey et al., 2021). Excessive intramural stress has been linked to inflammatory cell infiltration and transition of smooth muscle cells to a more macrophage-like phenotype (Li et al., 2020). Changes to mechanobiological stimuli are non-uniform down the PA tree in advanced disease, with values of WSS increasing in the distal vessels and decreasing in the proximal vessels (Ghorishi et al., 2007; Postles et al., 2014; Yang et al., 2019; Bartolo et al., 2022). This is likely driven by geometric differences in remodeling, where medial thickening causes distal narrowing and increased blood pressure proximally contributes to dilatation. However, the mechanisms underlying these differences in remodeling are poorly understood, and it is unknown how differences in baseline wall composition and mechanobiological environment may drive changes in different sized vessels of the PA tree.

Understanding differential drivers of vascular remodeling is further hindered by our inability to directly measure the mechanobiological stimuli experienced by cells. WSS and IMS are typically found using simulation tools that model the mechanical environment generated by hemodynamic loading (Marsden, 2013). Much work has been done on the simulation of hemodynamics in pulmonary hypertension to elucidate markers that predict disease progression and identify mechanobiological quantities of interest. Clinical imaging data has been used to build 3D models of the proximal vessels, and shown that increases in PVR are correlated with decreasing values of WSS in the proximal vessels (Kheyfets et al., 2015). WSS values were also found to increase along individual large vessel segments in healthy patients, but this trend was lost with the development of PAH (Zambrano et al., 2018).

For PAH, the mechanobiological stimuli of interest need to be quantified in vessels throughout the PA tree, including those too small to capture with standard medical imaging modalities. To overcome this, previous studies have used representative vascular networks, such as structured or morphometric trees, to estimate the loading environment in distal vessels (Qureshi et al., 2014; Yang et al., 2019; Bartolo et al., 2022), and varied the geometric properties of these networks with age and disease status (Dong et al., 2020; Dong et al., 2021). Changes in mechanical properties using linear models have been explored to capture the PA tree in hypoxic pulmonary hypertension (Acosta et al., 2017; Chambers et al., 2020). Mixture models that consider structurally significant constituents at homeostasis have been used to simulate the baseline PA tree (Gharahi et al., 2023). Additionally, models including the distal vasculature successfully predicted acute changes in blood flow distributions after PA interventions (Pries et al., 2001; Yang et al., 2016; Lan et al., 2022). Such models primarily relied on phenomenological rules to adapt blood vessel caliber in response to changes in loads. However, these adaptive networks were restricted to a healthy vessel phenotype, did not consider the PA tree as capable of elastic deformations, and did not attempt to model changes in vessel wall composition. Incorporating these effects becomes critical when considering models of disease progression in pathological vessels, as is the case in the setting of PAH. Geometric changes based on biological effects have also been used to change the PA tree morphometry to simulate disease progression in pulmonary hypertension, but these models do not simulate the time course of disease development (Postles et al., 2014).

Computational growth and remodeling (G&R) methods have been used to capture the evolution of single blood vessels and cardiovascular tissues. The constrained mixture theory G&R framework tracks the deposition and removal of structurally-significant constituents and solves for the changing loaded configuration of the vessel, making it well suited for predicting changes with altered loading and maladaptive biological stimuli (Szafron et al., 2018; Humphrey, 2021). Past works have described mechanobiological and immunological contributions to vessel remodeling and used numerical knockouts to identify pathways of importance for potential treatments (Latorre et al., 2020; Spronck et al., 2021). However, these models have not been used to simulate evolution in a vascular network and often use prescribed hemodynamic inputs rather than allowing for updates to pressure and flow based on changes in vascular resistance.

Herein, we develop a modeling framework to simulate the evolving hemodynamics of the entire PA tree in both adaptive and maladaptive remodeling cases. We apply a combination of morphometric trees and constrained mixture theory growth and remodeling to establish a loaded homeostatic state across different orders of the PA tree and add perturbations to baseline conditions that drive evolution of the hemodynamics over time. To understand the role of differences in behavior down the PA tree, we examine mechanobiological cues developed during adaptive remodeling and apply uniform and non-uniform stimuli to drive maladaptive remodeling. This framework represents an important step towards predicting the progression of PAH and evaluating the efficacy of therapies by using *in silico* studies.

## 2 Methods

### 2.1 Morphometric trees for hemodynamic simulations

Strahler diameter-defined morphometric trees with bifurcating branches were used to represent the pulmonary arterial vasculature of a mouse from the left pulmonary artery (LPA) down to the pre-capillary arteries (PrCAs) (Figure 1). Extensive details of morphometric trees have been outlined previously (Yang et al., 2019). These morphometric trees are generated from a connectivity matrix *C*_*mn*_, which gives the average number of vessel segments of order *m* that are connected to each segment of vessel order *n*, along with arrays of diameter *d*_*n*_ and length *l*_*n*_ values for each vessel order *n* ∈ [1, *N*]. The connectivity matrix from measurements made on the rat PA tree was used due to the lack of Strahler ordering data for the mouse (Jiang et al., 1994). To adapt rat data for generation of a murine tree, we removed the highest order entries of the rat tree connectivity matrix, which resulted in a tree with maximum order *N* = 10 for the LPA and a more appropriate total number of PrCAs with *n* = 1 for a mouse as determined by vessel counts from murine microCT (Townsley, 2012; Phillips et al., 2017). Boundary conditions for steady inlet flow and outlet pressure were specified from literature measurements of cardiac output (*CO*) and pulmonary capillary wedge pressure (*PCW*) in healthy adult mice, respectively, and mean pulmonary artery pressure (*mPAP*) was used to calculate the PA tree vascular resistance according to *PV R*_*meas*_ = (*mPAP* − *PCW*)*/CO* (Champion et al., 2000). The fractional flow split to the left lung *f*_*LP A*_ was taken from rat MRI data (Razavi et al., 2012). Parameter values for the homeostatic PA tree are included in Table 1.

**Figure 1:**
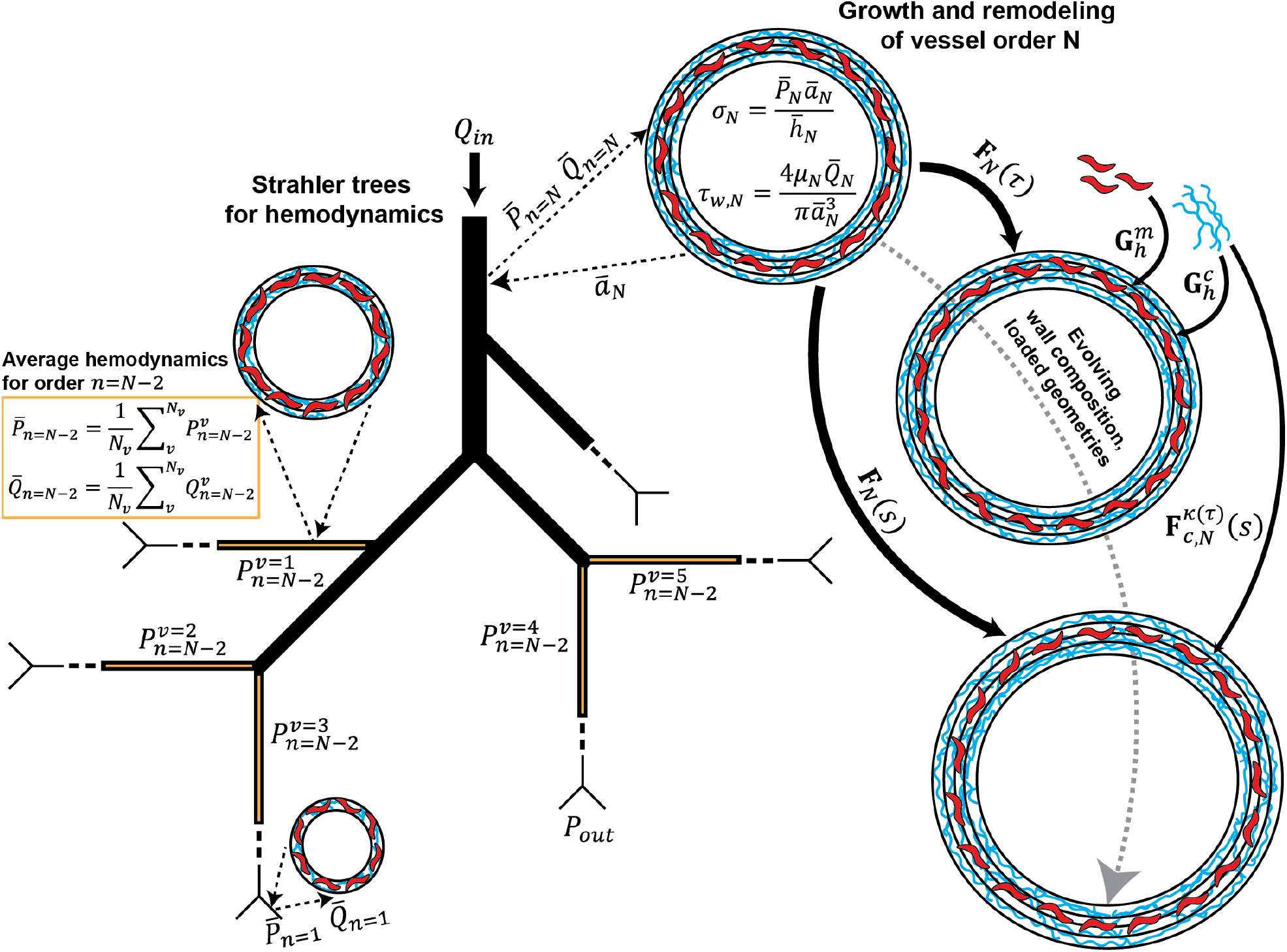
Schematic of morphometric tree-based framework for growth and remodeling of pulmonary arteries. Strahler diameter-defined morphometric trees are used to generate the bifurcating vessel tree (black lines, decreasing thickness for decreasing order). Inlet flow *Q*_*in*_ and outlet pressure *P*_*out*_ are prescribed as boundary conditions. For each vessel order, mean pressure 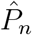 and mean flow 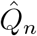 are passed to the G&R framework to determine the evolving average diameter *â*_*n*_. Examples of this exchange are shown for the orders *N, N* − 2, and 1. Mean values are passed for each order, shown here with cylinders representing vessels of decreasing size. Each vessel order can have its own composition, represented here by different relative amounts of smooth muscle (red), collagen (blue), and elastin (black) constituents. An example of mean hemodynamic calculation is shown for order *N* − 2, marked in orange, where values from all segments of that order are averaged to pass to the G&R framework. IMS *σ*_*N*_ and WSS *τ*_*w,N*_ are calculated in the G&R framework for each order. Configurations of vessel order *N* are shown over time to demonstrate the multiplicative combination of deformations that determines stretch of each constituent.

**Table 1:** Hemodynamic and morphometric parameters for generation of initial PA tree state

| Parameter | Description | Value | Units |
| --- | --- | --- | --- |
| $N$ | number of vessel orders | 10 | (-) |
| $d_N$ | LPA diameter<sup>1</sup> | 0.645 | (mm) |
| $l_N$ | LPA length<sup>1</sup> | 5.3 | (mm) |
| $h_N$ | LPA thickness<sup>1</sup> | 0.036 | (mm) |
| $f_d$ | order diameter ratio | 1.4 | (-) |
| $f_l$ | order length ratio<sup>2</sup> | 1.6 | (-) |
| $f_h$ | order thickness ratio | 0.06 | (-) |
| $mPAP$ | mean pulmonary arterial pressure<sup>3</sup> | 12.8 | (mmHg) |
| $PCW$ | pulmonary capillary wedge pressure<sup>3</sup> | 4.8 | (mmHg) |
| $CO$ | cardiac output<sup>3</sup> | 10.4 | (mL/min) |
| $f_{LPA}$ | LPA flow fraction<sup>4</sup> | 0.3 | (-) |

<sup>1</sup>Ramachandra et al., 2019
<sup>2</sup>Jiang et al., 1994
<sup>3</sup>Champion et al., 2000
<sup>4</sup>Razavi et al., 2012

Based on the observation of diameter and length scaling ratios, *f*_*d*_ and *f*_*l*_, which remain similar down the pulmonary arterial tree in various mammalian species (Huang et al., 1996), we identify order-specific diameters according to (*f*_*d*_)^*n*^*d*_*n*_ = *d*_*N*_ and order-specific lengths as (*f*_*l*_)^*n*^*l*_*n*_ = *l*_*N*_. Due to the greater dependence of hydraulic resistance on vessel diameter, we fix the length ratios to those of the rat, and optimize the diameter ratio *f*_*d*_, such that the difference in resistance of the morphometric tree from that derived with measured hemodynamic quantities is minimized according to the objective function *J*_*R*_ = (*PV R*_*calc*_ − *PV R*_*meas*_)^2^. Resistances for the morphometric tree are calculated by starting at the most distal branch and marching up the tree, where the resistance of a segment of order *n* is 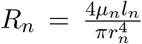 and resistance at each branch point is calculated using circuit rules for resistors in parallel as 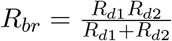 with *R*_*d*1_ and *R*_*d*2_ the resistances of each daughter branch. The viscosity *μ*_*n*_ is based on an empirical equation for microvessels in which the apparent viscosity is a function of diameter and discharge hematocrit due to the Fåhraeus-Lindqvist effect. (Pries et al., 1998; Secomb, 2017).

### 2.2 Homeostatic constrained mixture model of vessel mechanical behavior

Mechanical behavior of pulmonary arteries was considered as non-linear and anisotropic through application of a constrained mixture model. Previously published fits from biaxial extension and inflation experiments and histological evaluations of the C57BL/6 murine left pulmonary arteries were used to refit the material parameters for the homeostatic state of the highest order vessels in our model (*n* = 10) (Ramachandra et al., 2019). Passive stored energy density for the wall was considered as 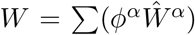 with *α* ∈ [*e*, m, c] for elastin, smooth muscle cells, and collagen, respectively. Homeostatic mass fractions *ϕ*^*α*^ were previously determined from histological evaluation. Constituent-specific bulk stored energy density functions 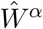 were prescribed for each of the different constituent types. For elastin, a neoHookean form was used with

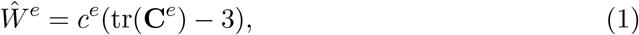

where *c*^*e*^ is the shear modulus and the elastin-specific right Cauchy-Green deformation tensor is **C**^*e*^ = (**F**^*e*^)^T^**F**^*e*^. The deformation gradient tensor for elastin is then **F**^*e*^ = **FG**^*e*^ with pre-stretch tensor 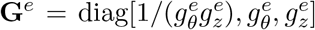 and 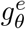 and 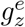 the circumferential and axial pre-stretch values. For the bulk stored energy density of smooth muscle cells, which are assumed to only bear load in the circumferential direction, an exponential, Fung-type form was used with

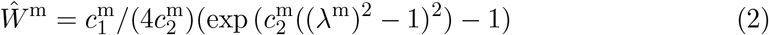

with 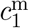 and 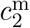 material parameters and *λ*^m^ = *λ*_*θ*_*g*^m^ with *g*^m^ the scalar pre-stretch of smooth muscle cells. For collagen, four distinct families *k* of fibers were considered, which contribute to load bearing in circumferential, axial, and diagonally symmetric directions. Each family is given a bulk stored energy density of

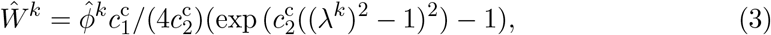

where 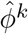 is the fraction of collagen fibers belonging to that family, 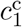 and 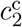 are material parameters common to all collagen fibers, and family-specific stretch in the fiber direction is 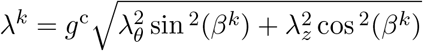 with *g*^c^ the collagen pre-stretch.

Cauchy stress can then be calculated constitutively as

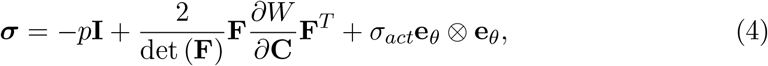

where Lagrange multiplier *p* enforces incompressibility and is determined by a plane stress assumption with *σ*_*rr*_ = 0 and active stress follows a Rachev-type contractility model with

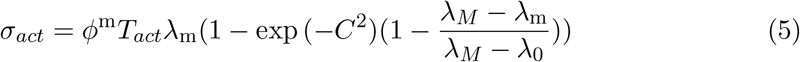

acting in the circumferential direction (Rachev et al., 1999). *ϕ*^m^ is the mass fraction of smooth muscle, *T*_*act*_ determines the magnitude of the active tone, *C* = *C*_*B*_ − *C*_*S*_Δ*τ*_*w*_ adapts tone in response to changes wall shear stress Δ*τ*_*w*_, which is zero for the homeostatic case, and *λ*_*M*_ and *λ*_0_ determine the stretch for peak and minimum activation, respectively. Circumferential and axial stresses from linear momentum balance for a thin walled cylinder are then 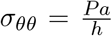 and 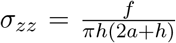, respectively, with *P* the transmural pressure, *a* the loaded inner radius, *h* the loaded thickness, and *f* the total axial force. Mechanical equilibrium was determined by balancing the constitutive and momentum balance forms for circumferential stress for a series of prescribed pressure and axial stretch states. Parameter values were found such that pre-stretch and active loading maintained an in vivo reference configuration, and a series of multiplicative deformations were used to simulate the experimental conditions with different loading states (Supplemental Figure 1). Least squares regression was used to find parameter values such that simulated stress vs strain results matched those found previously (Table 2), yielding excellent agreement between the current vessel modeling and the published PA behavior (Supplemental Figure 2).

**Table 2:** Baseline parameters for a constrained mixture model of the PA tree at homeostasis

| Parameters | Description | Values | Units |
| --- | --- | --- | --- |
| $\phi^e, \phi^m, \phi^c$ | constituent mass fractions | 0.255, 0.18, 0.565 | (-) |
| $c^e$ | elastin shear modulus | 40.73 | $kPa$ |
| $g_\theta^e, g_z^e$ | elastin pre-stretch | 1.11, 1.58 | (-) |
| $c_1^m, c_1^c$ | smooth muscle and collagen stiffness | 1.92, 87.22 | $kPa$ |
| $c_2^m, c_2^c$ | smooth muscle and collagen non-linearity | 0.16, 1.58 | (-) |
| $g^m, g^c$ | smooth muscle and collagen pre-stretch | 1.11, 1.11 | (-) |
| $\hat{\phi}^k$ | collagen fiber family fractions | [0.0 0.50 0.25 0.25] | (-) |
| $\beta^k$ | collagen fiber alignments | [90, 0, 26.32, -26.32] | $^\circ$ |
| $T_{act}$ | active stress magnitude | 36.13 | $kPa$ |
| $\lambda_M, \lambda_0$ | maximum and minimum active stretches | 1.1, 0.4 | (-) |
| $C_B, C_S$ | WSS modulation of active stress | 0.8326, 0.4163 | (-) |
| $\hat{k}^c, \hat{k}^T, \hat{k}^e$ | scaling parameters across each order | 0.0492, 0.895, 0.142 | (-) |
| $\hat{\phi}^T$ | minimal SMC contractility | 0.10 | (-) |
| $\hat{\rho}$ | mixture mass density | 1050 | $kg/m^3$ |
| $m_h^e, m_h^m, m_h^c$ | basal mass density production rates | 0.0, 42.15, 54.45 | $kg/(m^3 days)$ |
| $K_\sigma, K_\tau$ | mechanobiological gains for production | 4.0, 1.0 | (-) |
| $k_h^e, k_h^m, k_h^c$ | basal mass degradation rates | 0.0, 0.143, 0.143 | $days^{-1}$ |

### 2.3 Determining mechanical equilibrium of the pulmonary arterial tree

To analyze the entire pulmonary arterial tree, we calculated the mean luminal pressure at each vessel order from rigid hemodynamic simulations with the optimized diameter ratios. This average pressure was then used to determine the loaded geometry of all vessel segments of that order assuming a representative mechanical equilibrium problem for each order. Generally, in order *n* with vessel segments *v* ∈ [1, *N*_*v*_] and vessel specific pressures 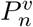, the mean pressure is calculated as

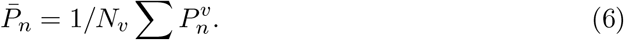

Then, mean Cauchy stress for each order is given by 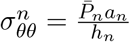. It has been shown that wall thickness varies down the PA tree, though the value of diameter to thickness ratio remains in a narrow range (Hislop et al., 1973). We therefore specify that order specific loaded thickness is given by *h*_*n*_ = *f*_*h*_*d*_*n*_ where *f*_*h*_ is a constant scaling ratio for thickness determined from the measured ratio in the LPA. As there is evidence that mechanical behavior changes with vessel size (Lee et al., 2016), we further allowed for the possibility of differing material parameters for different vessel orders by incorporating scaling ratios similar to those used for diameter. Homeostatic collagen mass fraction of each order was

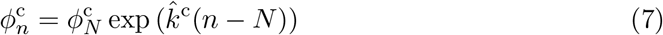

to allow for decreasing collagen content with adventitial thinning as vessel diameter decreases (Hislop et al., 1978). Basal values of the active stress parameter at each order were also allowed to change such that

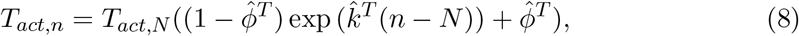

as there is evidence that PA smooth muscle contractility varies in different order vessels (Leach et al., 1992). 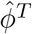 was included to maintain a minimal level of SMC contractility down the tree. It is also well known that elastin morphology changes between conduit and muscular arteries with a decrease in number of elastic lamellae and the presence of scattered elastic fibers within the media (Cocciolone et al., 2018). We therefore also allowed the elastin material behavior to change with vessel order according to

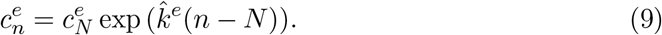

Values of scaling parameters 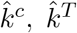, and 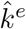 were optimized using non-linear regression to minimize an objective function that penalized differences between the prescribed intramural stress (from the geometry and pressure of the baseline tree) and that calculated constitutively assuming an *in vivo* reference configuration expressed as

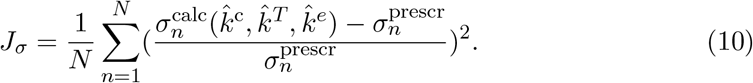

After the optimization converged, we again solved the equilibrium problem using average hemodynamic quantities to determine updated loaded geometries, and hemodynamic quantities were recalculated for the new geometry. Optimization of the material scaling ratios and updating of the hemodynamics was then repeated iteratively until the hemodynamics converged to a constant state. This converged homeostatic state and optimized material scaling ratios were then the basis for subsequent simulations that incorporated time course growth and remodeling. Optimized tree scaling parameter values are listed in Table 2.

### 2.4 Evolving constrained mixture model with time-varying composition and mechanical equilibrium

Additional equations are needed for the constrained mixture model to simulate the time course after a perturbation is made from the homeostatic state. The changing stored energy density of the mixture at time *s* is *W* (*s*) = ∑ *W* ^*α*^(*s*), where *W* ^*α*^(*s*) is the total stored energy density of each constituent *α* at time *s*. Following Valentin et al., 2009, we prescribe a form of

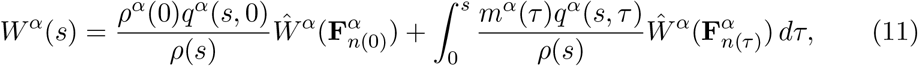

where *m*^*α*^(*s*) is the mass density production rate and *q*^*α*^(*s, τ*) is the survival fraction of the constituent produced at past time *τ* that survives to future time *s*. This form satisfies the mass fraction-based homeostatic case above and allows for cohorts of each constituent produced at past time *τ* to evolve their own deformation states at future times *s*. The evolving constituent specific deformation gradient is then 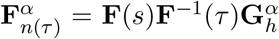 with mixture level deformation gradient **F**(*s*) and pre-stretch tensor 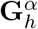 that is assumed here to remain constant with its homeostatic value. Mass densities of each constituent also change over time as

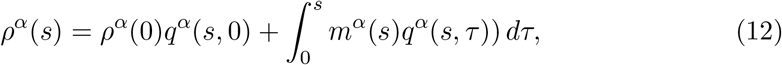

and the total mass density of the mixture can be found as *ρ*(*s*) = ∑*ρ*^*α*^(*s*). Note, both stored energy and mass densities are defined per unit reference volume. True mass density of the mixture 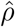 is assumed to remain constant, which allows for the volume change of the tissue to be calculated as det 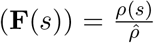. Solution of the mechanical equilibrium problem then proceeds as in the homeostatic case at each time step to determine the loaded radius and thickness.

For each constituent, three scalar equations, *m*^*α*^(*s*), *q*^*α*^(*s, τ*), and 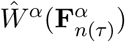 are required. Mass density production is defined as

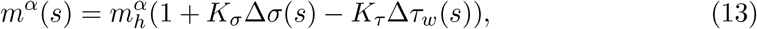

with 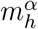 the homeostatic production rate. Mechanobiological stimuli modulate production with changes in intramural stress 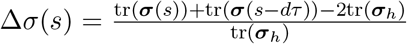 and wall shear stress 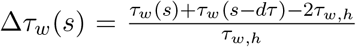 both affecting kinetics. These changes are calculated relative to the homeostatic values *σ*_*h*_ and *τ*_*w,h*_ for intramural and wall shear stress, which are set based on the loaded equilibrium state determined in initialization. As the hemodynamic coupling requires iteration at each fixed time point, incorporating an average of the mechanobiological stimuli from the previous time step *s* − *dτ* damps rapid changes and stabilizes the solution. Non-dimensional gains *K*_*σ*_ and *K*_*τ*_ control the magnitude of the response for all constituents to IMS and WSS, respectively. Survival functions for each constituent take the form 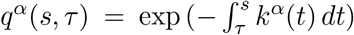 with degradation rate 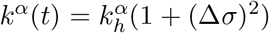 affected by changes in intramural stress from the homeostatic degradation rate 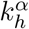. Functional forms for stored energy density 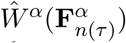 are the same as those used in the homeostatic case with a dependence on the constituent-specific, time varying deformation gradient 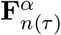. Parameter values for the growth and remodeling equations are listed in Table 2.

### 2.5 Novel constitutive relations for maladaptive remodeling

The baseline constrained mixture model framework accounts only for processes that are mechanobiologically-mediated, and are assumed to be adaptive. Past works simulating pathological remodeling also modified parameter values in specific relations or added additional terms that acted as a surrogate for biological stimuli in disease (Latorre et al., 2020; Spronck et al., 2021). Here, we outline a series of modifications to the baseline model that we used to build intuition for which modeling features are necessary to recapitulate key aspects of the clinical phenotypes observed in PAH.

To understand adaptive remodeling in response to flow changes, we modified the input flow according to

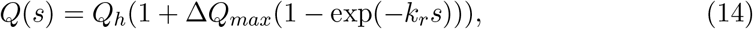

where Δ*Q*_*max*_ is the maximum change in flow from the homeostatic case *Q*_*h*_ and *k*_*r*_ is a rate parameter to modulate the rate of flow increase. Parametric studies with different values of Δ*Q*_*max*_ were performed to evaluate the ability of the coupled G&R and hemodynamic framework to simulate adaptation to stimuli similar to those experienced in PAH caused by congenital heart disease.

However, remodeling in PAH is often the consequence of maladaptive changes in vessel composition or geometry rather than from adaptive changes alone. We thus explored a series of changes to the model to account for maladaptation. To model hyperproliferation, we performed simulations with a modified form of the mass density production equations with

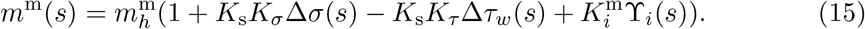

Here, 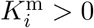 controls the magnitude of the increased proliferation and a stimulus function, such as 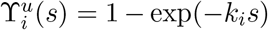 controls the time course. As PAH can be induced by increased WSS from a flow perturbation alone, we also performed simulations with an inflammatory stimulus based on the deviation in WSS from the homeostatic state according to a modified Hill function 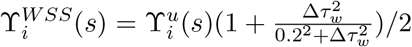 for Δ*τ* > 0. In PAH, normal responses to changes in mechanobiological stimuli are disrupted. We therefore included *K*_s_ ≤ 1 for all constituents as a modifier to the mechanobiological gains that reduces the sensitivity to the stimuli that allow for adaptive responses.

Functional forms for stored energy density of smooth muscle were also modified to allow for passive stiffening due to intracellular reorganization with changes in cell phenotype. We therefore prescribed

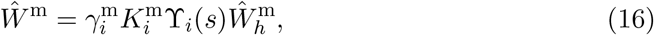

where stiffening occurs according to changes in 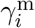 and the kinetics equation from mass density production. The phenotype of proliferating smooth muscle is often synthetic rather than contractile. Therefore, in maladaptation we explored an active stress form with

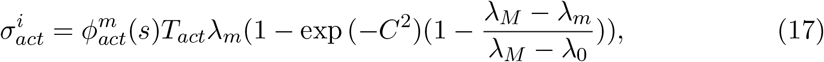

where the amount of contractile smooth muscle is fixed at the basal mass fraction with new proliferation assumed to be of a non-contractile phenotype using 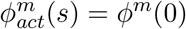. Finally, we allowed for loss of elastic fibers surrounding smooth muscle cells, as is often observed in PAH, by including

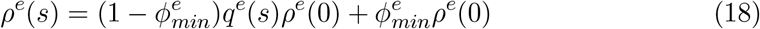

with a minimum persisting functional mass fraction of elastin 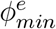 and degradation function *q*^*e*^(*s*) = *exp*(−*k*^*e*^*s*).

Different sets of these maladaptive stimuli were tested in cumulative combinations to determine their effects on geometric and hemodynamic evolution of the PA tree and identify those sets that led to physiologically realistic changes. In order, we added: hyperproliferation by including 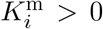 (Equation 15), passive stiffening of smooth muscle with 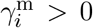 (Equation 16), loss of SMC contractility with 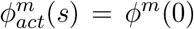 (Equation 17), loss of mechanobiological sensitivity *K*_s_ < 1 (Equation 15), and degradation of elastic fibers with 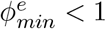 (Equation 18). Parameter values for simulations using each of the maladaptive modeling relations are listed in Table 3.

**Table 3:** Perturbation parameters for driving evolution of the PA tree

| Parameter | Description | Value | Units |
| --- | --- | --- | --- |
| $\Delta Q_{max}$ | fold increases in inlet flow | [0.0, 0.2, 0.4, 0.6, 0.8] | (-) |
| $k_r, k_i$ | rate of flow changes and proliferative onset | 0.25, 1.0 | $days^{-1}$ |
| $K_i^m$ | smooth muscle proliferative gain | 1.0 | (-) |
| $K_s$ | change in mechanobiological gain sensitivity | 0.25 | (-) |
| $\gamma_i^m$ | passive smooth muscle cell stiffening | 32 | (-) |
| $\phi_{min}^e$ | minimum elastin mass fraction | 0.5 | (-) |
| $k^e$ | elastin degradation rate | 0.0714 | $days^{-1}$ |

### 2.6 A coupled framework for evolving geometry and hemodynamics with varying perturbations

To simulate the evolution of the PA tree, we developed a coupled framework that passes pressure and flow information in each vessel order of the morphometric tree model to the constrained mixture-based G&R models, which then solves for the mechanical equilibrium state given those new hemodynamic inputs to update the tree geometry (Figure 1). The frameworks pass geometry and loads back and forth iteratively until the pressure solution converges at each time step, before continuing to march forward in time. At the homeostatic state, the system is stable with no change in geometry or hemodyanmics over time. Several different perturbations are then added to simulate different biological conditions. These perturbations are introduced sequentially with different parameters of the model modified to understand the interactions of each constituent and process.

## 3 Results

### 3.1 Varying material properties down the PA tree are needed to maintain homeostatic geometry

We first simulated pressure in the PA tree with prescribed values of radii, which decrease with decreasing vessel order (Figure 2A), matching the PVR of the simulated tree to literature measurements on mice (Figure 2B). We then simulated pressure in the loaded state with constant material parameters from fitting the LPA biaxial mechanical data and using pressure values from the prescribed geometry, i.e. setting 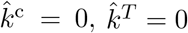, and 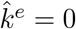. The resulting loaded configuration diameters for constant material parameters down the PA tree were then used to update the hemodynamics. Mean pulmonary arterial pressures were more than 3-fold higher than the values found with the prescribed rigid geometry matched to literature values (Champion et al., 2000), illustrating that uniform parameters do not allow for a physiologic loading state of the PA tree (Figure 2B). As such, values for the scaling parameters 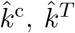, and 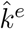 were fit to minimize the difference of the loaded tree from the prescribed rigid tree. With the optimized parameter set, we obtained less than 5% error in loaded vs. prescribed radius measurement (Figure 2C), and pressures and radii at each order of the tree were similar to those estimated for the rigid tree (Figure 2B and C).

**Figure 2:**
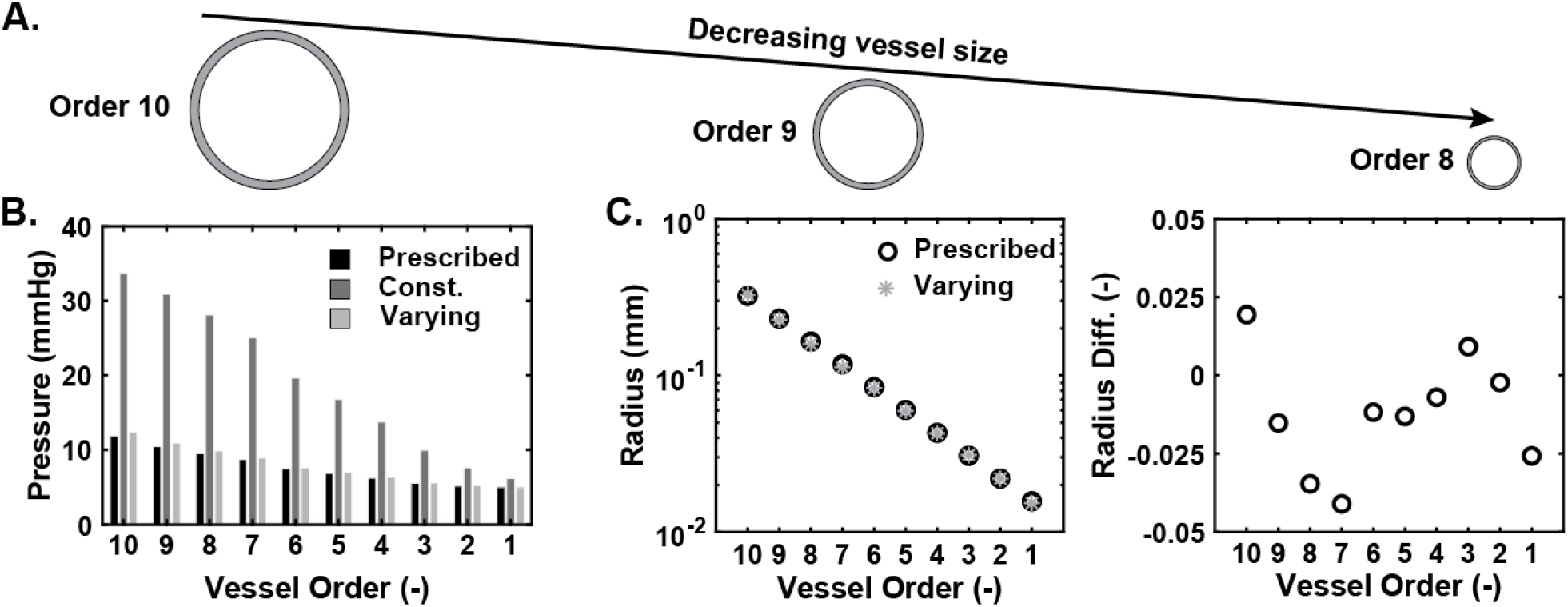
Homeostatic pressure and radius down the PA tree. **A)** Schematic illustrating the decrease in diameter as vessel order decreases with relative vessel sizes to scale. **B)** Pressure down the PA tree for each vessel order with radii values prescribed to match the literature value of PVR, radius values found after pre-loading with constant material parameters, and radius values found after pre-loading with variable material behavior down the tree fit to the prescribed state. **C)** Radius comparisons for the prescribed and varying cases (left) and the relative difference in radius after fitting (right).

For the optimized scaling parameters, collagen content was found to decrease by ∼20% down the PA tree (Figure 2A). Smooth muscle and elastin content were assumed to equally increase as collagen content decreased, leading to ∼10% increases of each. Despite a decrease in the elastin material parameter of 75% from the highest to lowest orders, we observed structural stiffening with decreasing vessel order, as normalized increases in pressure from the homeostatic pressure in each order led to smaller changes in radius from their baseline value (Figure 3B). The basal active stress parameter also decreased with vessel order, approaching the asymptotic lower threshold by order 4 with vessel diameters on the order of 80 *μ*m (Figure 3C).

**Figure 3:**
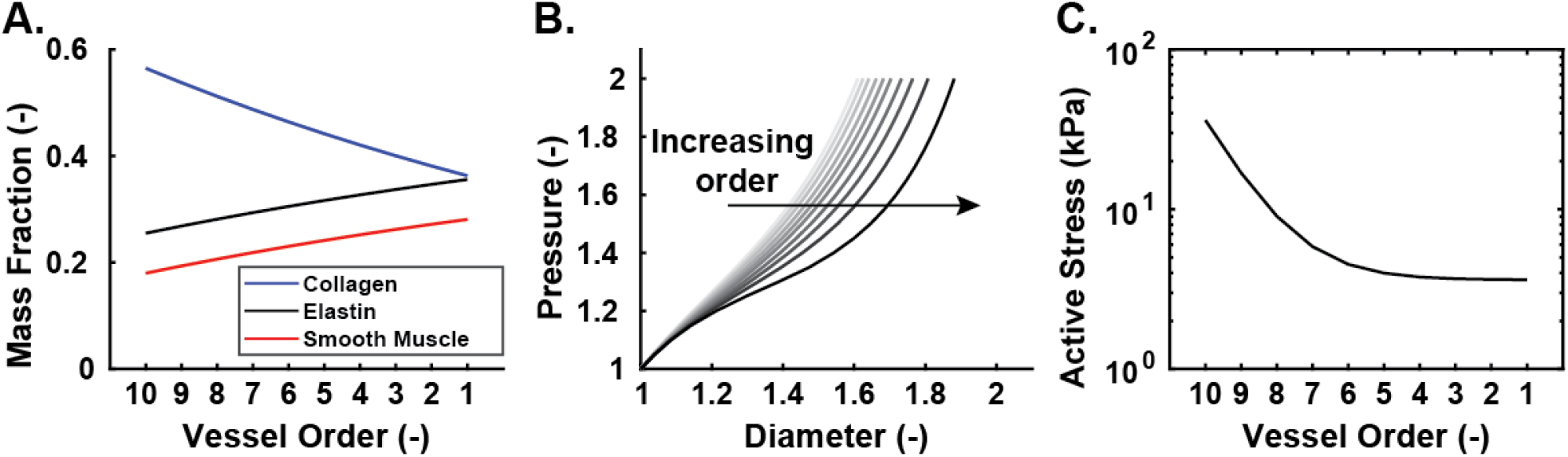
Differences in homeostatic composition and behavior for each order of the PA tree. **A)** Mass fractions of collagen, elastin, and smooth muscle cells for each vessel order. **B)** Pressure vs. diameter behavior for each vessel order with each step normalized by its order-specific homeostatic value. **C)** Active stress parameter values for each vessel order.

### 3.2 Hemodynamic feedback is essential for modeling PA tree adaptation

We next evaluated the need for hemodynamic feedback, i.e. the passing of updated flow and pressure information as geometry changes at each time step of the growth and remodeling time course. First, we tested a case with a 40% increase in inlet flow, simulating a mild left-to-right shunt as found in congenital heart disease patients who may develop PAH, and computed the corresponding change in pressure and flow at each order without allowing for any change in geometry from the homeostatic case, i.e. *ā*_*n*_ was not returned to the morphometric tree for iterative updates to the hemodynamics. Thus, pressures in the large vessels increased greatly relative to the homeostatic case, as increases in small artery diameter to adapt to increased WSS did not lower PVR and decrease pressure. In response to high pressures, higher order vessels dilated and thickened without coming to a new equilibrium state (Figure 4A). IMS increased sharply in the larger vessels, while gradual increases were seen in the small vessels as they dilated to adapt to increased flow. WSS decreased in most orders beyond its initial value as the dilatation from increased pressurization outweighed the contribution to WSS from the increased flow.

**Figure 4:**
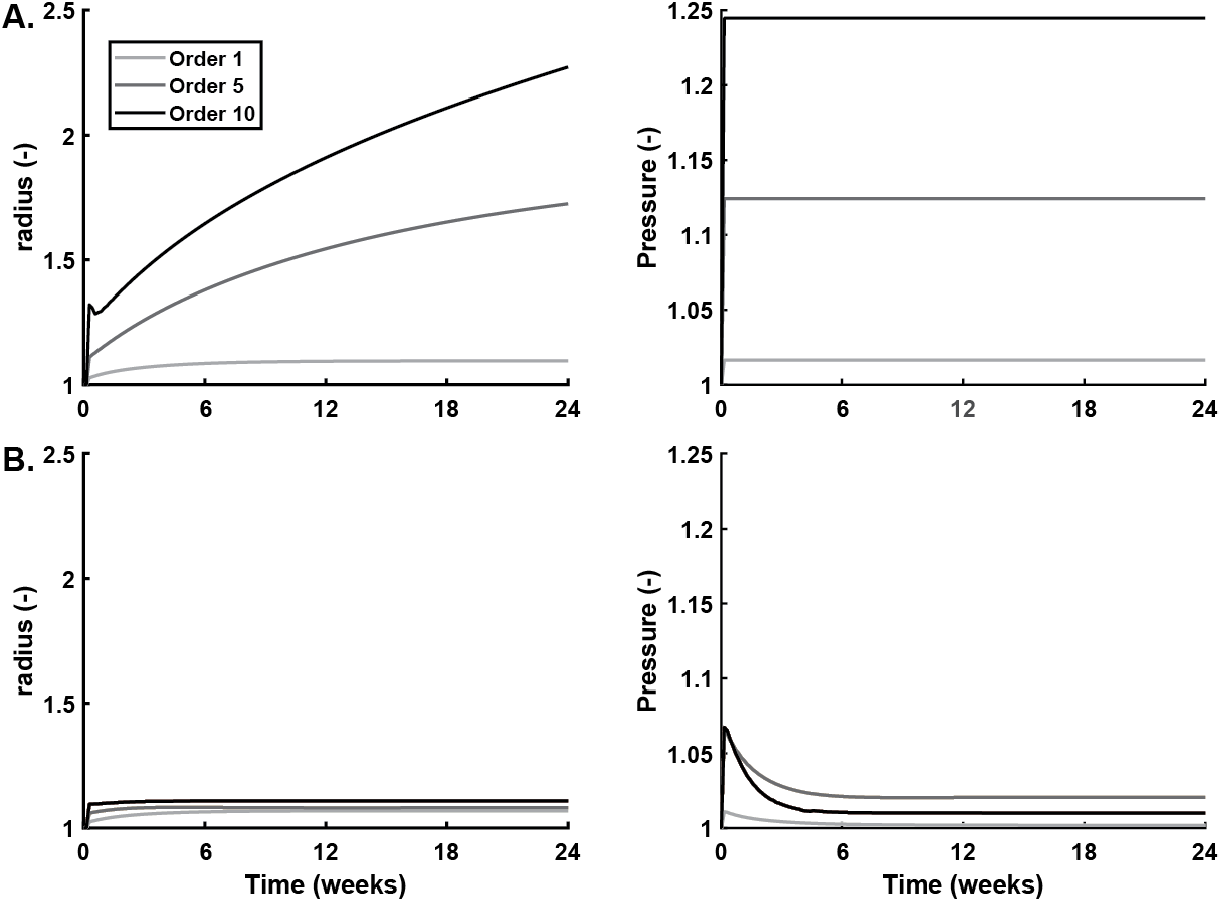
Effects of including hemodynamic feedback after a flow perturbation. Remodeling of radius and pressure for cases both without **A)** and with **B)** hemodynamic feedback after a 40% increase in inlet flow. Initial pressures after the flow perturbation are used for the entire G&R time course in the case without feedback. Values are normalized by those at the homeostatic state.

These changes are not physiologic, as pressure only begins to increase 4 weeks after an increase in flow in a murine model of PAH induced by an abdominal aortocaval shunt (Zhang et al., 2018). Thus, we retained the contribution of hemodynamic feedback in the model for the remainder of the study, where changes in geometry were used to recalculate the pressure and flow at each point during G&R. The initial pressure increase was therefore blunted by elastic dilatation of all vessels after the perturbation, and subsequently decreased with dilatation of all vessels in the PA tree (Figure 4B). Thickness of all vessels decreased slightly to allow for the dilatation. IMS and WSS were both elevated in all vessel orders.

### 3.3 Adaptive remodeling in response to flow decreases PVR and maintains vasodilatory capacity

To further understand the effects of increasing flow on PA tree adaptation, we performed a parametric study with increasing inlet flow perturbations from 5% to 80%, as would be expected with ventricular septal defects of small to moderate sizes (Dong et al., 2021). In all cases and for all vessel orders, the pulmonary arteries dilated and wall shear stress increased (Figure 5). Progressive increases in inlet flow led to more dilatation and greater increases in wall shear stress. Dilatation increased for larger vessels with Order 10 vessels dilating more than Order 5 or Order 1 vessels. This corresponded to larger changes in wall shear stress from the homeostatic state with a nearly 4-fold relative increase in wall shear stress in Order 1 vessels in comparison to Order 10 vessels. Furthermore, larger vessels more quickly reestablished an equilibrium state, with Order 10 vessels only taking 3 weeks to reach a new steady-state (defined by less than 0.05% change in radius between time steps), while Order 1 vessels took around 6 weeks. Intramural stress also plays a key mechanobiological role in G&R, which is affected by changes in thickness. In Order 10 vessels, thickness changes were modest, while decreases in thickness were observed for Order 5 and Order 1 vessels (Supplemental Figure 3). Thickness was observed to decrease to a greater degree as the inlet flow was increased. The decrease in thickness along with increase in radius led to increases in intramural stress with increasing inlet flow and with decreasing vessel size.

**Figure 5:**
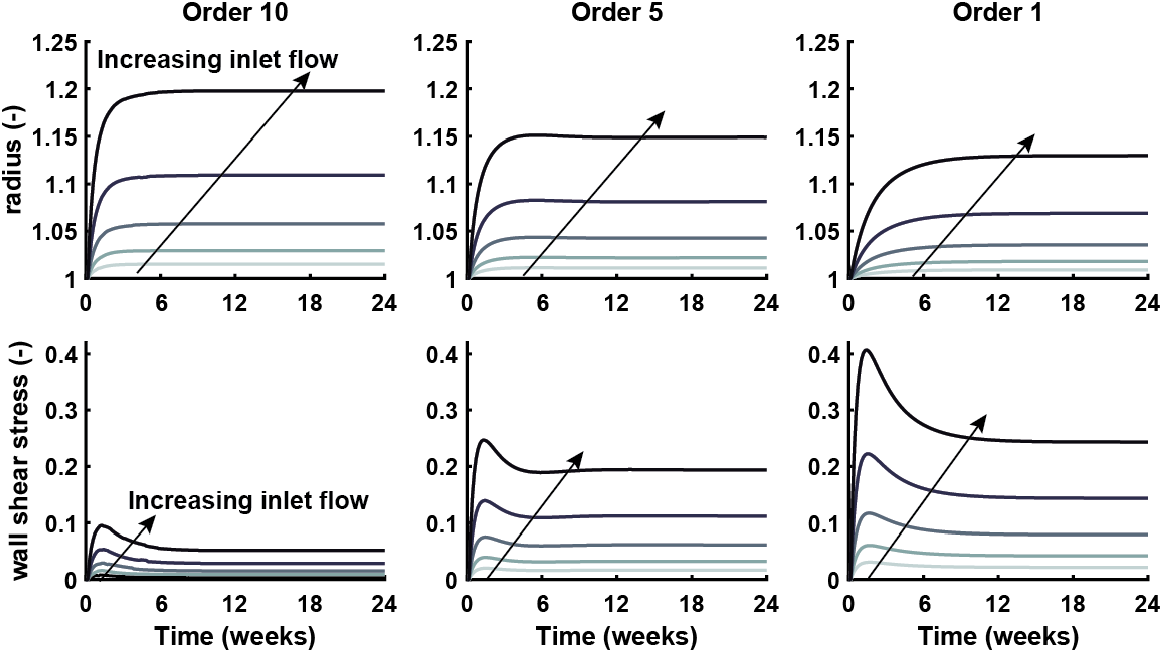
Evolution of PA tree with increasing inlet flow. Changes in radius and wall shear stress over time after 5, 10, 20, 40, and 80% increases in flow, i.e. Δ*Q*_*max*_ = [0.05, 0.1, 0.2, 0.4, 0.8], for vessel orders 10, 5 and 1. Values are normalized by their homeostatic state.

We also evaluated pressure evolution in the largest order vessels of the PA tree for each of the flow cases and modeled pulmonary vascular resistance changes with decreasing active tone in smooth muscle cells to simulate the effects of vasodilator administration (Figure 6). Pressures increased with increasing flow rates but remained near their homeostatic levels, increasing less than 2% at the steady-state for an 80% increase in flow (Figure 6A). Baseline PVR with normal active tone decreased with increasing flow (Figure 6B), as would be necessary to maintain the nearly normal afterload on the right ventricle (left). The fold change in PVR was similar across all flow changes (right), indicating that smooth muscle function is not compromised by adaptive remodeling.

**Figure 6:**
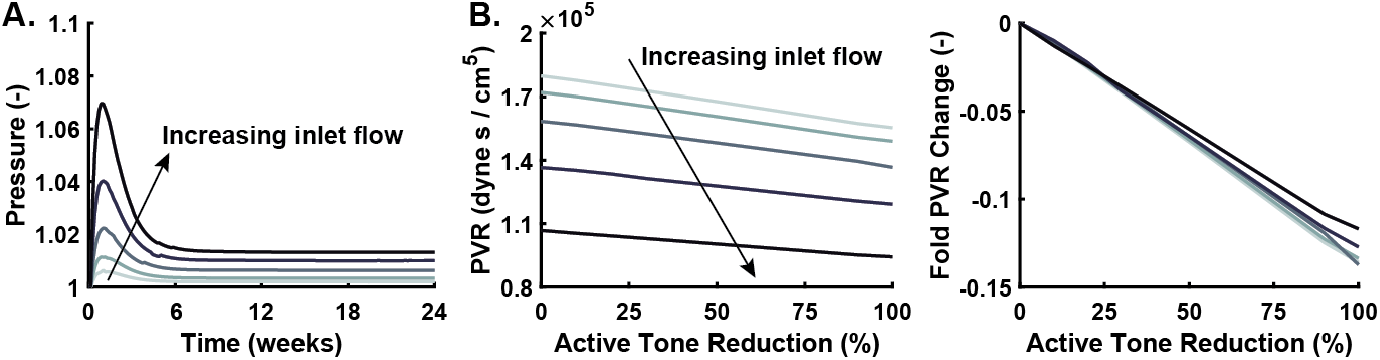
Pressure and resistance changes with increasing inlet flow. **A)** Change in pressure of Order 10 vessels corresponding to the LPA over time for each of the flow perturbations in Figure 5. Values are normalized by their homeostatic state. **B)** Change in pulmonary vascular resistance (PVR) with reduction of active smooth muscle tone after 24 weeks of remodeling for each of the flow perturbations (left). Fold changes in PVR are also shown from the state with no reduction of active tone (right).

### 3.4 Uniform maladaptive stimuli applied across vessel orders

We next determined what combination of maladaptive stimuli across vessel orders could lead to pathophysiologic geometric changes, namely dilatation of the high order, proximal vessels, and narrowing of the low order, distal vessels with corresponding decreases and increases in wall shear stress, respectively (Kheyfets et al., 2015; Yang et al., 2019). Maladaptive stimuli were imposed using the mathematical relations outlined in Section **2.5** with each successive stimuli applied cumulatively (Figure 7). For prescribed hyperplasia of smooth muscle cells, the functional form 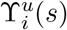 was used. Parameter values for each simulation are included in Table 3. Addition of SMC hyperplasia alone caused proximal vessel narrowing and distal vessel dilatation, while SMC hyperplasia in conjunction with passive stiffening led to narrowing of all vessel orders. Removing the contractile capacity of new smooth muscle cells showed limited effects in lower order vessels, while higher order vessels had less narrowing and nearly maintained their original radius. Reduction of mechanobiological sensitivity in combination with all prior stimuli led to some dilatation and decrease of WSS in the large order vessels, while the small order vessels narrowed and had substantial elevation of WSS. Further addition of elastin degradation to the cumulative stimuli showed narrowing of large order vessels and effects on distal vessels were modest. We similarly examined the effects of these cumulative stimuli on thickness and intramural stress (Supplemental Figure 4). SMC hyperplasia alone had led to a slight decrease in thickness and intramural stress for all orders. When combined with passive SMC stiffening, thickness decreases were more pronounced in the smaller order vessels but minimal in the highest order vessels, and increases in intramural stress were caused by increased pressure. The change to a synthetic SMC phenotype for new proliferation only affected the largest order vessels and led to a slight increase in radius and intramural stress. Mechanobiological dysfunction led to much larger increases in thickness for the large and mid-sized vessels, and consequently decreased intramural stresses. Elastin loss tempered the thickness increases in the large and mid-sized vessels, and further decreased thickness of the smallest order vessels.

**Figure 7:**
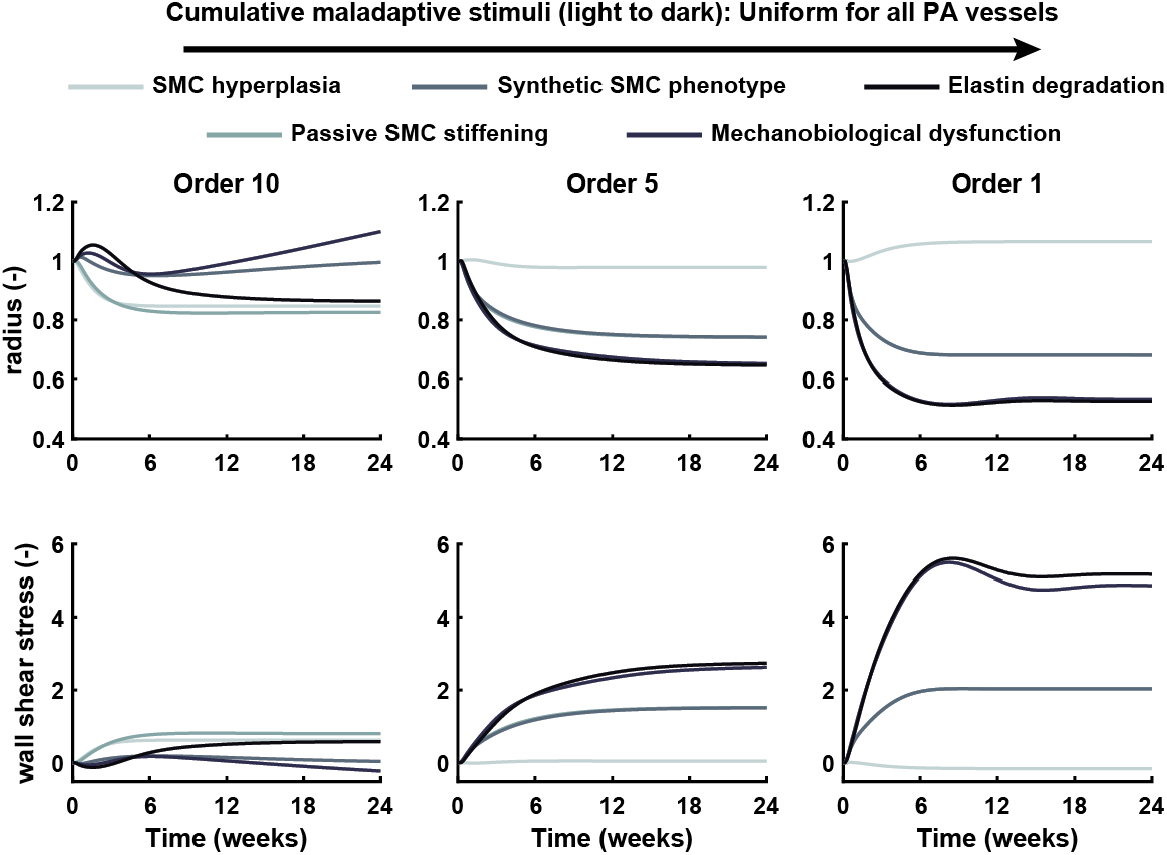
Uniform maladaptive stimuli applied throughout the PA tree. Maldaptive stimuli applied cumulatively. Each darker-colored line includes the effects of stimuli from the simulation results plotted with the lighter-colored lines. Changes in radius and wall shear stress over time for vessel orders 10, 5 and 1 were found with SMC hyperplasia 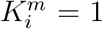, passive stiffening of SMCs 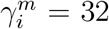, a synthetic SMC phenotype for proliferating cells 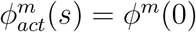, a decrease in normal mechanobiological function *K*_s_ = 0.25, and elastin degradation to 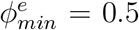. Values of radius and wall shear stress are normalized by those from their homeostatic state.

Changes in pressure evolution in the largest order vessels of the PA tree and in pulmonary vascular resistance with decreasing active tone were modeled for the combinations of maladaptive stimuli (Figure 8). With increased smooth muscle proliferation alone, pressure increased modestly (Figure 8A). For each of the other cases, pressure increased drastically with a 2- to 4-fold increase in pressure. Increased smooth muscle proliferation alone increased PVR by 10%, but allowed for much greater reduction of PVR with reduction of active tone. In all other cases, pulmonary vascular resistance increased at least 3-fold, and the ability to reduce PVR with decreasing active tone was almost completely lost; none of the cases were able to reduce PVR more than 5%.

**Figure 8:**
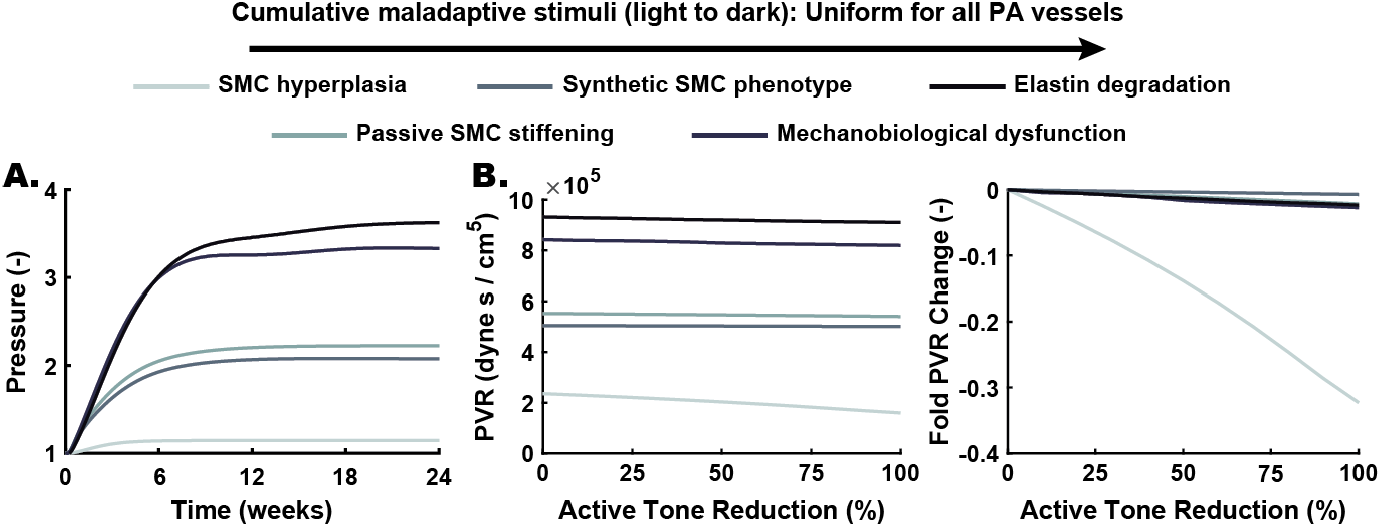
Pressure and resistance changes with uniform maladaptive stimuli. Maladaptive stimuli applied cumulatively as in Figure 7. Each darker-colored line includes the effects of stimuli from the simulation results plotted with the lighter-colored lines. **A)** Change in pressure of Order 10 vessels corresponding to the LPA over time for corresponding stimuli in Figure 7. Values are normalized by their homeostatic state. **B)** Change in pulmonary vascular resistance (PVR) with reduction of active smooth muscle tone after 24 weeks of remodeling for each of the uniform maladaptive stimuli combinations (left). Fold changes in PVR are also shown from the state with no reduction of active tone (right).

### 3.5 Wall shear stress-based maladaptaion drives strongly non-uniform remodeling

Due to the contribution of altered WSS to changes in endothelial phenotype, we next trialed a form for a maladaptive stimulus function driven by increased WSS, 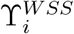, as outlined in **Section 2.5**. This was applied in conjunction with other maladaptive stimuli applied uniformly across the tree, consisting of increased passive SMC stiffness, decreased mechanosensitivity, and a transition from contractile to synthetic phenotype for new SMC production with the parameter values from the prior case. The development of PAH was then instigated by the application of an 80% increase in flow from baseline with Δ*Q*_*max*_ = 0.8. As in the case with all uniform maladaptive stimuli, dilatation occurred in the largest order vessels and narrowing was evident in the smallest order vessels (Figure 9). However, the degree of dilatation was much greater, as the largest order vessels had diameters at least 3 times higher than their original values. A sharp transition was observed between Order 7 and Order 6 vessels partway down the PA tree, where Order 6 vessels narrowed modestly, and Order 7 vessels dilated significantly. This was caused by the transition from elevated WSS to decreased WSS at Order 7 and higher vessels, which did not induce as much SMC proliferation. WSS values decreased greatly for the higher order vessels that dilated and increased in the smallest order vessels up to 4 times higher than their initial values. Further addition of elastin degradation had minimal impact on the resulting vessel evolution.

**Figure 9:**
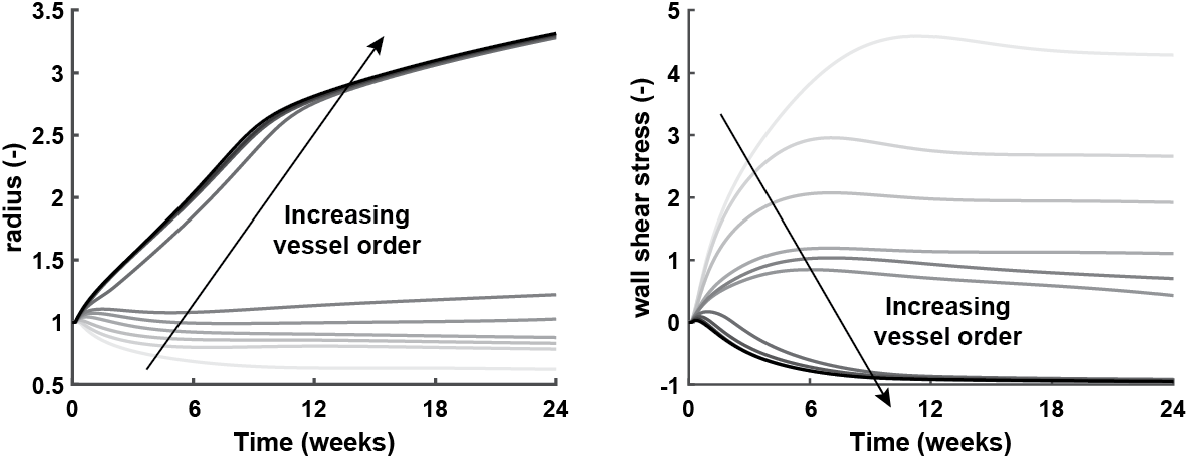
Wall shear stress-mediated inflammation leads to non-uniform PA remodeling. Evolution after an 80% increase in flow with Δ*Q*_*max*_ = 0.8 for radius (left) and WSS (right) with maladaptive remodeling from SMC proliferation mediated by WSS according to 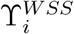. Uniform maladaptive stimuli were applied for passive SMC stiffening, loss of mechanosensitivity, and loss of a contractile SMC phenotype as in the prior case. Values are normalized to their homeostatic state and darkening lines show remodeling for vessels of increasing caliber.

## 4 Discussion

Disease progression in pulmonary arterial hypertension is driven by remodeling of the PA vasculature. In this work, we developed a framework for coupled simulations of hemodynamics, wall mechanics, and composition of the pulmonary arterial tree during the development and progression of PAH. This framework allows for the imposition of maladaptive remodeling stimuli with uniform or spatially varying behavior, thus enabling the identification of mechanisms that most contribute to increasing disease severity. Importantly, it also can be used to predict the time course of critical metrics of clinical interest, such as the PAP, PVR, and potential responsiveness to vasodilatation. Our modeling approach allowed for novel consideration of elastic effects on PA tree hemodynamics. When establishing the homeostatic loaded configuration of our PA tree, we confirmed the critical need for varying material and contractile behavior for vessels of different sizes (Leach et al., 1992), as uniform properties yielded non-physiologic hemodynamics (Figure 2). Compositional changes identified by our model after fitting the loaded state of the PA tree to match the homeostatic PVR agree with literature data on changes in amounts of collagen for smaller vessels (Figure 3), as distal vessels have relatively thinner adventitia while maintaining a similar overall wall thickness relative to lumen size (Townsley, 2012). Previous experiments have also identified increasing structural stiffness of the PA tree with decreasing vessel diameter (Lee et al., 2016), which are consistent with our predictions of decreasing compliance with decreasing caliber. Furthermore, we demonstrated the need for hemodynamic feedback between the G&R framework and the fluid mechanics solution for the PA tree (Figure 4), as the changes in diameter and thickness from altered mechanobiological stimuli can allow for adaptation to perturbations in loading conditions. Without this feedback, pulmonary arterial pressure increases immediately when there is an increase in flow, which does not agree with literature demonstrating a capacity for the PA tree to maintain afterload for weeks after this type of insult (Zhang et al., 2018).

There is a delay in the initiation of PAH after a flow increase in animal models, where PAP increases 4 weeks after an aorto-caval shunt is created in a mouse model (Zhang et al., 2018). In an ovine model of PAH, different degrees of flow insults cause varying disease severity, and assays of cell function and distal wall morphometry show modest differences in cases with less severe disease despite significant changes to hemodynamics (Kameny et al., 2019). Using our modeling framework, we show that mechanobiological cues that drive vascular remodeling deviate from their homeostatic state even after allowing for a period of adaptive remodeling (Figure 5, Supplemental Figure 3). Deviations in WSS and IMS were found to be greater in the distal vessels whose narrowing causes increased PVR and PAP. This suggests that despite the initial adaptability of the PA tree, long-term exposure to abnormal mechanobiological stimuli of sufficient magnitude can initiate PAH through the eventual transition from adaptive to maladaptive responses by mechanosensitive cells in the distal vessels, which then raises pressures and subsequently alters proximal hemodynamic loading. Early measurements of PAP and PVR do not reflect this abnormal mechanobiological environment in situations with increased flow, as PAP remains near its normal value and PVR initially decreases as the PA tree adapts to increasing flow (Figure 6). However, as the disease progresses these adaptive processes are no longer able to compensate for the abnormal hemodynamic environment. Distal remodeling triggered by persistent elevation in WSS eventually contributes to vessel narrowing, which leads to a positive feedback loop of rising WSS over time in the distal vessels that drives the maladaptive remodeling of vessels throughout the PA tree. With this narrowing of distal vessels, PVR increases and leads to end-stage disease with right ventricular failure and death. At the point when adaptive remodeling has failed to restore the normal mechanobiological environment, there are numerous biological pathways activated as part of the maladaptive remodeling response, including vascular cell dysfunction, inflammatory cell infiltration, and alteration of the extracellular matrix (Humbert et al., 2008; Rabinovitch et al., 2014; Tojais et al., 2017). Smooth muscle cells effect many of these changes, and were thus the focus of our initial studies on developing constitutive relations that contribute to vascular changes akin to those seen in patients with PAH. We added maladaptive stimuli cumulatively to find the combination that led to distal narrowing and proximal dilatation as observed clinically (Yang et al., 2019). At minimum, increased smooth muscle proliferation and passive stiffening of smooth muscle was required to instigate distal narrowing when applied uniformly through the PA tree, though proximal vessels also narrowed in contrast to clinical observation (Figure 7). Only with additional disruption to normal SMC behavior, in which case adaptive contractility and mechanosensitivity was lost, did the proximal vessels dilate. This highlights the complex interactions between pathways that contribute to PAH, as considering only activation of inflammatory-like responses, i.e. proliferation and stiffening, did not capture the clinically observed behavior. Furthermore, loss of elastin mass from all vessel orders led to non-physiologic remodeling despite much documentation of elastin damage in PAH, suggesting that careful selection of mathematical relations is necessary for matching model behavior to observed results. Effects of elastin damage could also play a biochemical role that could be captured through phenotypic changes of cells and deposition of other matrix constituents. Indeed, identifying such deviations from expected behavior with a prescribed insult can provide new directions for experiments, such as a need for isolating biomechanical and signaling effects of elastin degradation. Thickening observed in the proximal vessels was greater than that observed experimentally (Supplemental Figure 4), suggesting that additional tuning of parameter values and functional forms may be required to fully recapitulate all aspects of the clinical PAH phenotype (Prapa et al., 2013).

Different cases of cumulative stimuli added here led to remodeling similar to both moderate and severe cases of PAH as determined by the simulated changes in pressure (Figure 8). As such, there is a need for clinical simulation tools that reflect the mechanobiological environment of each patient to better inform the potential for disease progression. Current methods allow for simulation of WSS at single time points for a patient-specific model (Tang et al., 2012; Yang et al., 2019), but future coupling of this G&R framework with 3D computational fluid dynamics simulations will offer improved tools for interpreting and predicting the stimuli that contribute to PAH development. At different times in disease progression, varying combinations of maladaptive stimuli may be needed to more closely match the continued development of PAH. Thus,preventing further deterioration in a patient’s condition could require targeting of different pathways at different times depending on disease severity. By validating this model formulation against time course data on development of PAH, future work should focus on identifying critical times and pathways for successful intervention as well as determining critical junctures when remodeling becomes irreversible.

The diverse etiologies of PAH likely mean that multiple combinations of maladaptive stimuli with different functional forms are required to model progression in realistic disease scenarios. For flow-induced PAH, as in the case of left-to-right shunt from a ventricular septal defect, high WSS is thought to play a critical role in disease initiation. We therefore modified the constitutive relation for SMC proliferation to depend on increased WSS from the homeostatic value, which led to non-uniform remodeling through the PA tree and a moderate PAH phentoype caused by distal vessel narrowing with increased WSS (Figure 9). However, low and oscillatory WSS can also contribute to endothelial dysfunction, like that commonly found in atherosclerotic plaques (Deng et al., 2021). Proximal vessels experience low WSS after dilatation, which could activate similar remodeling pathways as in distal vessels and act to homogenize the effective behavior of the PA tree despite very different values of mechanobiological stimuli in different orders of the PA. Our future modeling efforts will therefore also examine the differences in uniform vs. non-uniform stimuli needed to match time course changes in experimental models of PAH.

There are several limitations to our approach. Homogenization of G&R for all vessels of each order, where average hemodynamics determine the geometric and compositional changes, greatly simplifies the computations necessary for predictions of disease progression but decreases the fidelity of simulated spatial variations for vessels of the same size that are in different parts of the PA tree. Some of those vessels with greatly decreased flow after narrowing may also undergo pruning, which is not currently considered in this formulation, as the morphometry of the tree remains fixed at all time points. Hemodynamics for each blood vessel are calculated using Poiseuille flow assumptions with a fluid circuit analogy, which is less accurate than 1D or 3D solutions of the Navier-Stokes equations. 1D solutions have been used previously in the distal vessels, which also allows better examination of unsteady flow profiles (Bartolo et al., 2022). Metrics like oscillatory shear index that come from time-varying flows could also be a significant mechanobiological factor in disease progression, as has been seen in the systemic circulation (Yang et al., 2018). Finally, many parameter values are assumed from the literature and require future experimental validation, along with confirmation of appropriate functional forms for maladaptive stimuli that capture time course remodeling data. However, the approach demonstrated herein was chosen to allow for simulations with both adaptive and maladaptive remodeling stimuli, which have not been performed previously. These assumptions allow for an efficient solution process, where prediction of a single time course for the PA tree can take less than 15 minutes on a laptop computer. Modifying any of these assumptions would greatly increase the cost, as there are several iterative steps to ensuring convergence of the model with passing of hemodynamic and geometric variables from the solution of flow equations in the morphometric tree to the G&R equations for the wall mechanics. Trialing different functional forms for maladaptive stimuli is then prohibitively expensive with much longer compute times. To remedy these limitations, models with different orders of fidelity should be developed, such that many runs of simulations with lower fidelity models can inform the use of higher fidelity simulations.

This model represents an important step towards predicting the evolution of PAH. Outputs of this model are of direct clinical interest, and the ability to tune specific parameters that relate to biological pathways gives our framework the unique ability to be used for simulating interventions. With further validation, the model could be used to test promising interventions or treatment scenarios that target specific stimuli or pathways, ultimately leading to improved outcomes for PAH patients.

## 7 Supplemental Figures

**Supplemental Figure 1:**
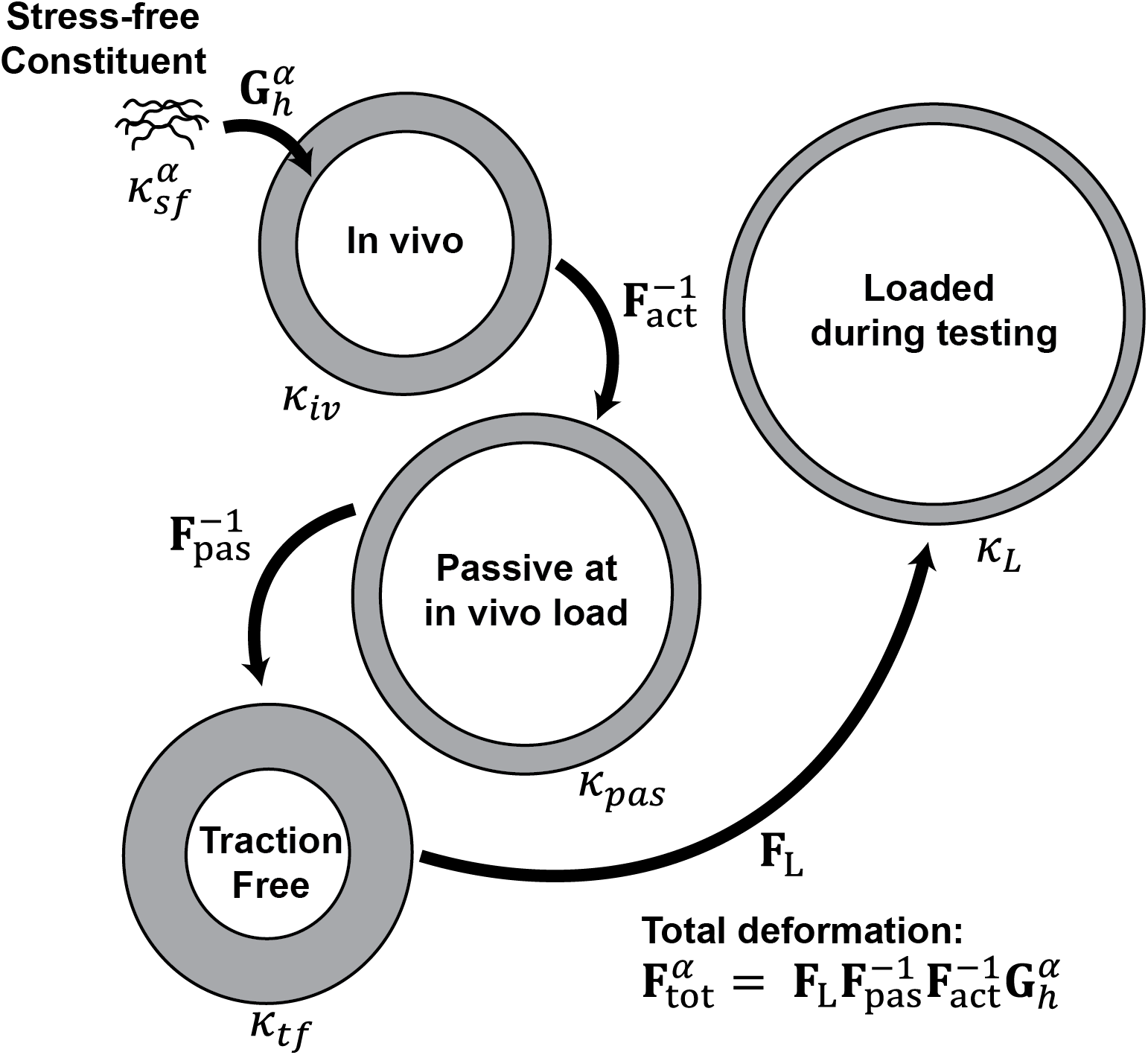
Configurations and deformations for capturing LPA mechanical behavior. To fit the pre-stretch of each constituent 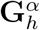, we considered the the total deformation experienced by each constituent during mechanical testing 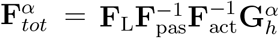. Constituents were assumed to be deposited from their stress-free natural configuration 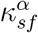 into the in vivo configuration *κ*_*iv*_ with pre-stretch 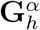. The biaxial mechanical testing was performed in the passive state, therefore deformation 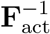 was prescribed from active contraction data to set the passive loaded configuration *κ*_*pas*_. Then, deformation 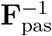 was found for the traction-free configuration *κ*_*tf*_. Then, the loaded configuration at each loaded state was found with deformation **F**_L_ to the loaded configuration *κ*_*L*_ for a specific pressure and axial stretch.

**Supplemental Figure 2:**
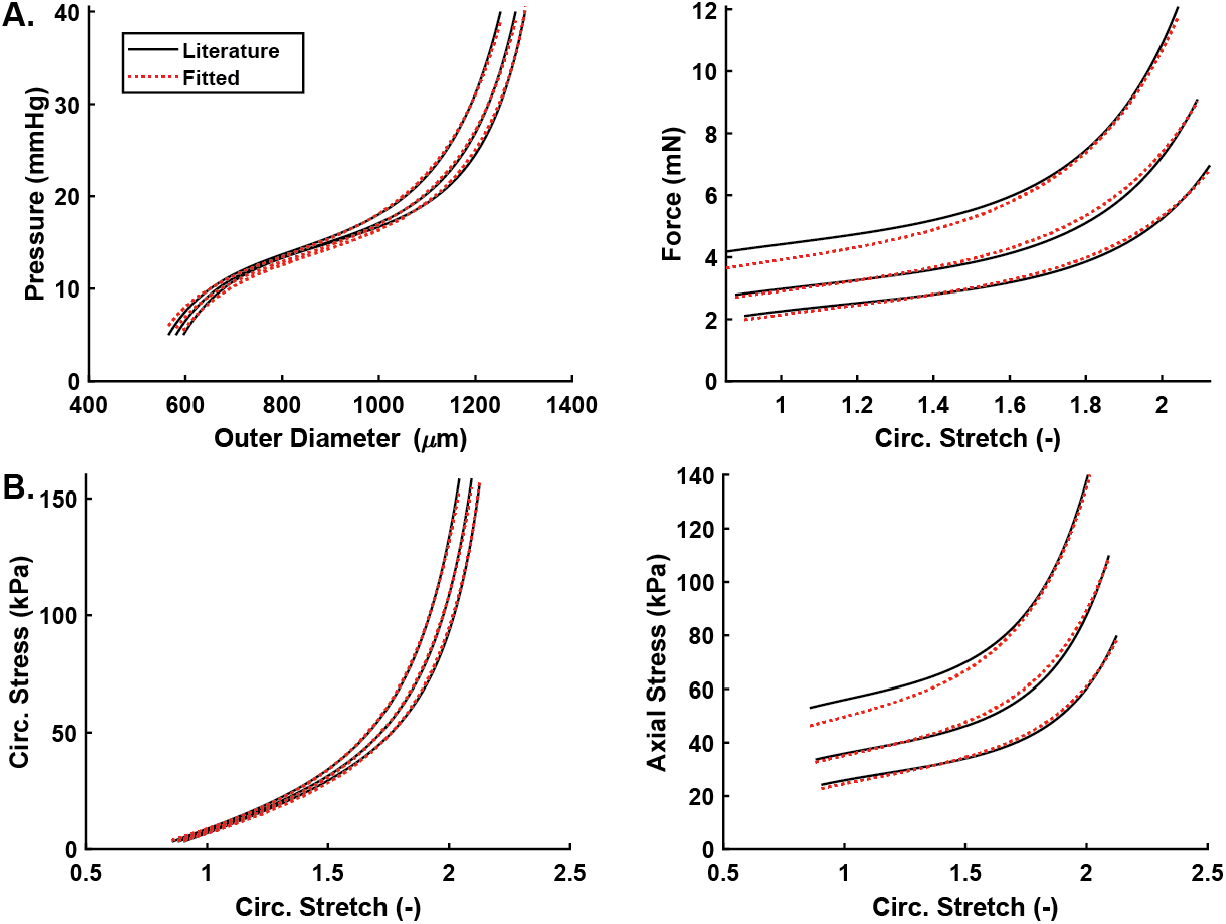
Fitted behavior for constrained mixture model of LPA behavior. **A)** Pressure vs. diameter behavior and Axial force vs. circumferential stretch for 3 tests at the in vivo value of axial stretch and values -5% and +5% with the literature results from Ramachandra and Humphrey, 2019 (black) and the fitted results with the constrained mixture model (red dashed) (Ramachandra et al., 2019. **B)** Circumferential stress vs. strain behavior using the literature parameter values and those fitted with the constrained mixture model.

**Supplemental Figure 3:**
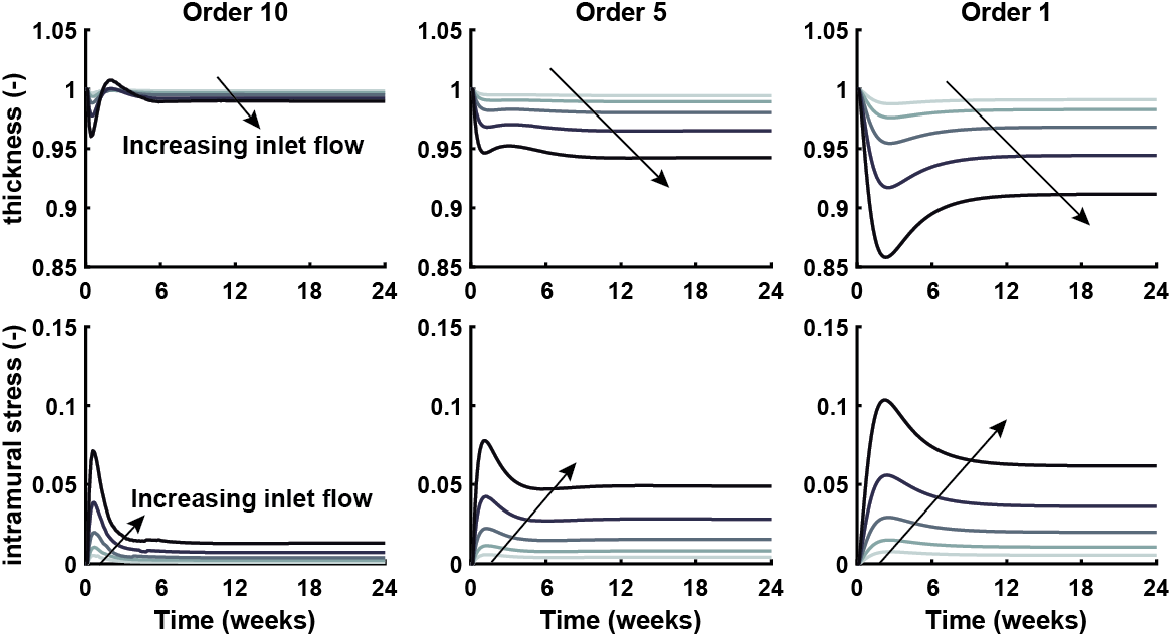
Evolution of thickness and intramural stress for increasing inlet flow. Changes in thickness and intramural stress over time after 5, 10, 20, 40, and 80% increases in flow, i.e. *dQ* = [0.05, 0.1, 0.2, 0.4, 0.8], for vessel orders 10, 5 and 1. Values are normalized by their homeostatic state.

**Supplemental Figure 4:**
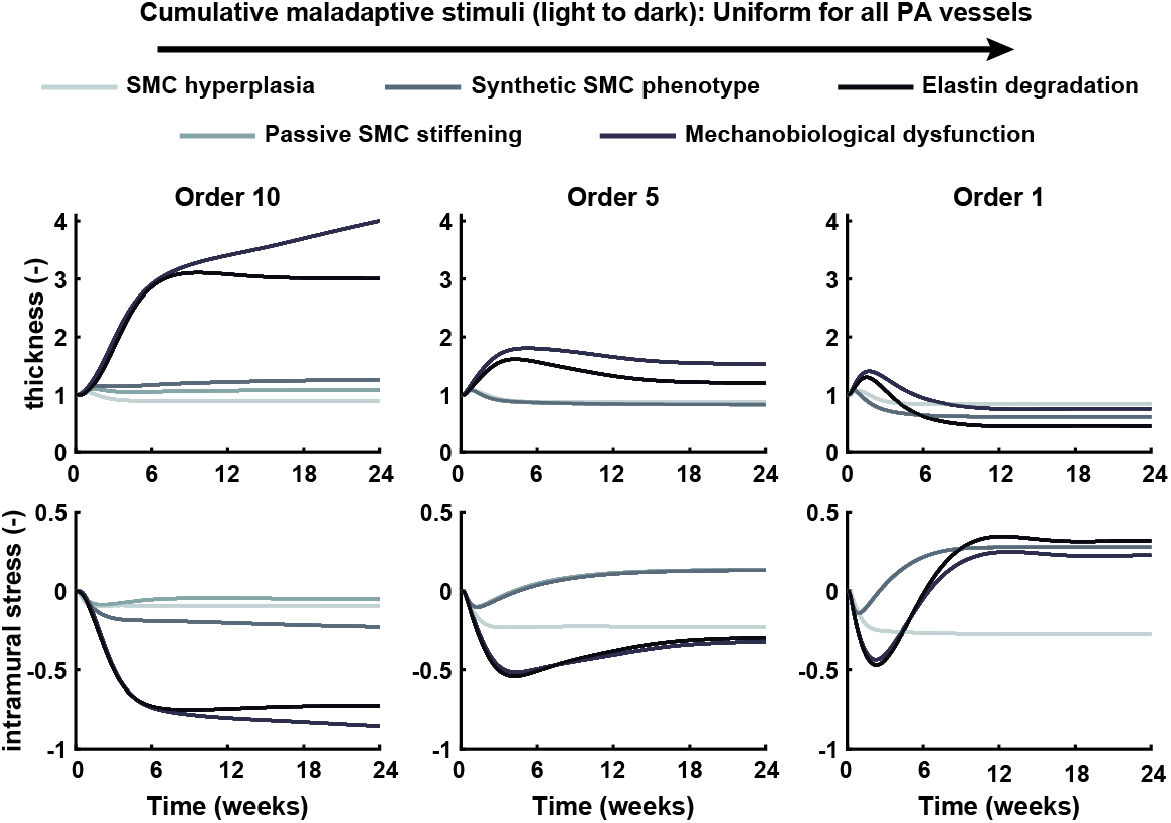
Thickness and intramural stress with uniform maladaptive stimuli applied throughout the PA tree. Maladaptive stimuli applied cumulatively. Each darker-colored line includes the effects of stimuli from the simulation results plotted with the lighter-colored lines. Changes in thickness and intramural stress over time for vessel orders 10, 5 and 1 were found with SMC hyperplasia 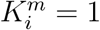, passive stiffening of SMCs 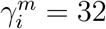, a synthetic SMC phenotype for proliferating cells 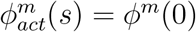, a decrease in normal mechanobiological function *K*_s_ = 0.25, and elastin degradation to 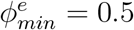. Values of thickness and intramural stress are normalized by their homeostatic state.

## 8 Acknowledgements

The authors appreciate funding support from the Parker B. Francis Fellowship program for this work. This work was also supported in part by by the Stanford Maternal and Child Health Research Institute through the Pilot Award Program and a grant from the National Heart, Lung and Blood Institute (1T32HL098049).

## Notes

### Competing Interest Statement

The authors have declared no competing interest.

## References

Abman, Steven H et al. (2015). “Pediatric pulmonary hypertension: guidelines from the American heart association and American thoracic Society”. In: Circulation 132.21, pp. 2037–2099.

Acosta, Sebastián et al. (2017). “Cardiovascular mechanics in the early stages of pulmonary hypertension: a computational study”. In: Biomechanics and modeling in mechanobiology 16, pp. 2093–2112.

Bartolo, Michelle A et al. (2022). “Numerical predictions of shear stress and cyclic stretch in pulmonary hypertension due to left heart failure”. In: Biomechanics and Modeling in Mechanobiology 21.1, pp. 363–381.

Chambers, Megan J et al. (2020). “Structural and hemodynamic properties of murine pulmonary arterial networks under hypoxia-induced pulmonary hypertension”. In: Proceedings of the Institution of Mechanical Engineers, Part H: Journal of Engineering in Medicine 234.11, pp. 1312–1329.

Champion, Hunter C et al. (2000). “A novel right-heart catheterization technique for in vivo measurement of vascular responses in lungs of intact mice”. In: American Journal of Physiology-Heart and Circulatory Physiology 278.1, H8–H15.

Cocciolone, Austin J et al. (2018). “Elastin, arterial mechanics, and cardiovascular disease”. In: American Journal of Physiology-Heart and Circulatory Physiology 315.2, H189–H205.

Deng, Hanqiang et al. (2021). “Activation of Smad2/3 signaling by low fluid shear stress mediates artery inward remodeling”. In: Proceedings of the National Academy of Sciences 118.37, e2105339118.

Dong, Melody et al. (2020). “Image-based scaling laws for somatic growth and pulmonary artery morphometry from infancy to adulthood”. In: American Journal of Physiology-Heart and Circulatory Physiology 319.2, H432–H442.

Dong, Melody L et al. (2021). “Computational simulation-derived hemodynamic and biomechanical properties of the pulmonary arterial tree early in the course of ventricular septal defects”. In: Biomechanics and Modeling in Mechanobiology 20.6, pp. 2471–2489.

Gharahi, Hamidreza et al. (2023). “A multiscale framework for defining homeostasis in distal vascular trees: applications to the pulmonary circulation”. In: Biomechanics and Modeling in Mechanobiology, pp. 1–16.

Ghorishi, Zahra et al. (2007). “Shear stress paradigm for perinatal fractal arterial network remodeling in lambs with pulmonary hypertension and increased pulmonary blood flow”. In: American Journal of Physiology-Heart and Circulatory Physiology 292.6, H3006–H3018.

Hislop, A and L Reid (1978). “Normal structure and dimensions of the pulmonary arteries in the rat.” In: Journal of anatomy 125.Pt 1, p. 71.

Hislop, Alison and Lynne Reid (1973). “Pulmonary arterial development during childhood: branching pattern and structure”. In: Thorax 28.2, pp. 129–135.

Huang, Wei et al. (1996). “Morphometry of the human pulmonary vasculature”. In: Journal of applied physiology 81.5, pp. 2123–2133.

Humbert, Marc et al. (2008). “Endothelial cell dysfunction and cross talk between endothelium and smooth muscle cells in pulmonary arterial hypertension”. In: Vascular pharmacology 49.4-6, pp. 113–118.

Humphrey, Jay D and Martin A Schwartz (2021). “Vascular mechanobiology: homeostasis, adaptation, and disease”. In: Annual Review of Biomedical Engineering 23, pp. 1–27.

Humphrey, JD (2021). “Constrained mixture models of soft tissue growth and remodeling–twenty years after”. In: Journal of Elasticity 145.1, pp. 49–75.

Ivy, Dunbar (2016). “Pulmonary hypertension in children”. In: Cardiology clinics 34.3, pp. 451–472.

Jiang, ZL, GS Kassab, and YC Fung (1994). “Diameter-defined Strahler system and connectivity matrix of the pulmonary arterial tree”. In: Journal of Applied Physiology 76.2, pp. 882–892.

Kameny, Rebecca Johnson et al. (2019). “Ovine models of congenital heart disease and the consequences of hemodynamic alterations for pulmonary artery remodeling”. In: American journal of respiratory cell and molecular biology 60.5, pp. 503–514.

Kheyfets, Vitaly O et al. (2015). “Patient-specific computational modeling of blood flow in the pulmonary arterial circulation”. In: Computer methods and programs in biomedicine 120.2, pp. 88–101.

Lan, Ingrid S et al. (2022). “Virtual Transcatheter Interventions for Peripheral Pulmonary Artery Stenosis in Williams and Alagille Syndromes”. In: Journal of the American Heart Association 11.6, e023532.

Latorre, Marcos and Jay D Humphrey (2020). “Numerical knockouts–In silico assessment of factors predisposing to thoracic aortic aneurysms”. In: PLoS computational biology 16.10, e1008273.

Leach, RM et al. (1992). “A comparison of the pharmacological and mechanical properties in vitro of large and small pulmonary arteries of the rat”. In: Clinical science (London, England: 1979) 82.1, pp. 55–62.

Lee, Pilhwa et al. (2016). “Heterogeneous mechanics of the mouse pulmonary arterial network”. In: Biomechanics and modeling in mechanobiology 15.5, pp. 1245–1261.

Li, Guangxin et al. (2020). “Chronic mTOR activation induces a degradative smooth muscle cell phenotype”. In: The Journal of clinical investigation 130.3, pp. 1233–1251.

Marsden, Alison L (2013). “Simulation based planning of surgical interventions in pediatric cardiology”. In: Physics of fluids 25.10, p. 101303.

Moonen, Jan-Renier et al. (2015). “Endothelial-to-mesenchymal transition contributes to fibro-proliferative vascular disease and is modulated by fluid shear stress”. In: Cardiovascular research 108.3, pp. 377–386.

Moonen, Jan-Renier et al. (2022). “KLF4 recruits SWI/SNF to increase chromatin accessibility and reprogram the endothelial enhancer landscape under laminar shear stress”. In: Nature communications 13.1, pp. 1–16.

Phillips, Michael R et al. (2017). “A method for evaluating the murine pulmonary vasculature using micro-computed tomography”. In: Journal of Surgical Research 207, pp. 115–122.

Postles, Arthur, Alys R Clark, and Merryn H Tawhai (2014). “Dynamic blood flow and wall shear stress in pulmonary hypertensive disease”. In: 2014 36th annual international conference of the IEEE engineering in medicine and biology society. IEEE, pp. 5671–5674.

Prapa, Matina et al. (2013). “Histopathology of the great vessels in patients with pulmonary arterial hypertension in association with congenital heart disease: large pulmonary arteries matter too”. In: International journal of cardiology 168.3, pp. 2248–2254.

Pries, AR, B Reglin, and TW Secomb (2001). “Structural adaptation of microvascular networks: functional roles of adaptive responses”. In: American Journal of Physiology-Heart and Circulatory Physiology 281.3, H1015–H1025.

Pries, AR, TW Secomb, and P Gaehtgens (1998). “Structural adaptation and stability of microvascular networks: theory and simulations”. In: American Journal of Physiology-Heart and Circulatory Physiology 275.2, H349–H360.

Qureshi, M Umar et al. (2014). “Numerical simulation of blood flow and pressure drop in the pulmonary arterial and venous circulation”. In: Biomechanics and modeling in mechanobiology 13.5, pp. 1137–1154.

Rabinovitch, Marlene et al. (2014). “Inflammation and immunity in the pathogenesis of pulmonary arterial hypertension”. In: Circulation research 115.1, pp. 165–175.

Rachev, Alexander and Kozaburo Hayashi (1999). “Theoretical study of the effects of vascular smooth muscle contraction on strain and stress distributions in arteries”. In: Annals of biomedical engineering 27.4, pp. 459–468.

Ramachandra, Abhay B and Jay D Humphrey (2019). “Biomechanical characterization of murine pulmonary arteries”. In: Journal of biomechanics 84, pp. 18–26.

Razavi, Hedi et al. (2012). “A method for quantitative characterization of growth in the 3-D structure of rat pulmonary arteries”. In: Microvascular research 83.2, pp. 146–153.

Secomb, Timothy W (2017). “Blood flow in the microcirculation”. In: Annual Review of Fluid Mechanics 49, pp. 443–461.

Spronck, Bart et al. (2021). “Excessive adventitial stress drives inflammation-mediated fibrosis in hypertensive aortic remodelling in mice”. In: Journal of the Royal Society Interface 18.180, p. 20210336.

Szafron, JM et al. (2018). “Immuno-driven and mechano-mediated neotissue formation in tissue engineered vascular grafts”. In: Annals of biomedical engineering 46.11, pp. 1938–1950.

Tang, Beverly T et al. (2012). “Wall shear stress is decreased in the pulmonary arteries of patients with pulmonary arterial hypertension: an image-based, computational fluid dynamics study”. In: Pulmonary circulation 2.4, pp. 470–476.

Tojais, Nancy F et al. (2017). “Codependence of bone morphogenetic protein receptor 2 and transforming growth factor-β in elastic fiber assembly and its perturbation in pulmonary arterial hypertension”. In: Arteriosclerosis, thrombosis, and vascular biology 37.8, pp. 1559–1569.

Townsley, Mary I (2012). “Structure and composition of pulmonary arteries, capillaries and veins”. In: Comprehensive Physiology 2, p. 675.

Valentin, A et al. (2009). “Complementary vasoactivity and matrix remodelling in arterial adaptations to altered flow and pressure”. In: Journal of The Royal Society Interface 6.32, pp. 293–306.

Yang, Tung-Lin et al. (2018). “Differential regulations of fibronectin and laminin in Smad2 activation in vascular endothelial cells in response to disturbed flow”. In: Journal of Biomedical Science 25.1, pp. 1–13.

Yang, Weiguang, Jeffrey A Feinstein, and Irene E Vignon-Clementel (2016). “Adaptive outflow boundary conditions improve post-operative predictions after repair of peripheral pulmonary artery stenosis”. In: Biomechanics and modeling in mechanobiology 15, pp. 1345–1353.

Yang, Weiguang et al. (2019). “Evolution of hemodynamic forces in the pulmonary tree with progressively worsening pulmonary arterial hypertension in pediatric patients”. In: Biomechanics and modeling in mechanobiology 18.3, pp. 779–796.

Zambrano, Byron A et al. (2018). “Image-based computational assessment of vascular wall mechanics and hemodynamics in pulmonary arterial hypertension patients”. In: Journal of biomechanics 68, pp. 84–92.

Zhang, Mingjie et al. (2018). “Characteristics of pulmonary vascular remodeling in a novel model of shunt-associated pulmonary arterial hypertension”. In: Medical Science Monitor: International Medical Journal of Experimental and Clinical Research 24, p. 1624.

